# Glyphosate Effects on Growth and Biofilm Formation in Bee Gut Symbionts and Diverse Associated Bacteria

**DOI:** 10.1101/2024.03.20.585985

**Authors:** Erick V. S. Motta, Tyler K. de Jong, Alejandra Gage, Joseph A. Edwards, Nancy A. Moran

## Abstract

Biofilm formation is a common adaptation enabling bacteria to thrive in various environments and to withstand external pressures. In the context of host-microbe interactions, biofilms play vital roles in establishing microbiomes associated with animals and plants and are used by opportunistic microbes to facilitate proliferation within hosts. Investigating biofilm dynamics, composition, and responses to environmental stressors is crucial for understanding microbial community assembly and biofilm regulation in health and disease. In this study, we explore the independent gut colonization and in vitro biofilm formation abilities of core members of the honey bee (*Apis mellifera*) gut microbiota. Additionally, we assess the impact of glyphosate, a widely used herbicide with antimicrobial properties, and a glyphosate-based formulation on growth and biofilm formation in bee gut symbionts as well as in other biofilm-forming bacteria associated with diverse animals and plants. Our results demonstrate that several strains of core bee gut bacterial species can independently colonize the bee gut, which probably depends on their ability to form biofilms. Furthermore, glyphosate exposure has varying effects on bacterial growth and biofilm formation. These findings imply specific impacts of environmental stressors on microbial biofilms with both ecological and host health-related implications.

Importance

Biofilms are essential for microbial communities to establish and thrive in diverse environments. In the honey bee gut, the core microbiota member *Snodgrassella alvi* forms biofilms, potentially aiding the establishment of other members and promoting interactions with the host. In this study, we show that specific strains of other core members, including *Bifidobacterium*, *Bombilactobacillus*, *Gilliamella*, and *Lactobacillus*, also form biofilms. We then examine the impact of glyphosate, a widely used herbicide that disrupts the bee microbiota, on their growth and biofilm formation. Our findings demonstrate diverse effects of glyphosate on biofilm formation, ranging from inhibition to enhancement, reflecting observations in other beneficial or pathogenic bacteria associated with animals and plants. Thus, glyphosate exposure may influence bacterial growth and biofilm formation, potentially shaping microbial establishment on host surfaces and impacting health outcomes.

## Introduction

Biofilms are complex structures formed by microorganisms that stably adhere to surfaces by building a structured matrix of extracellular polymeric substances (1–4). They provide protection to microbes against antimicrobial agents (5, 6) and other hostile conditions (7), and also facilitate communication (8, 9), recombination and gene transfer (10), and nutrient exchange within the community (11, 12).

Biofilm development can occur on diverse surfaces, both natural and artificial (13). In interactions with animals and plants, microbial biofilms are found in various body compartments, playing crucial roles in health and disease (14). For instance, in humans, gut microbial biofilms aid in nutrient metabolism, immune system regulation, and competitive exclusion of pathogens (8, 15). In plants, microbial biofilms contribute significantly to nutrient cycling, plant growth, and pathogen protection in the rhizosphere and phyllosphere (16–18). However, some opportunistic microbes, such as *Pseudomonas aeruginosa*, form biofilms on animal and plant surfaces and thereby proliferate and cause disease (14, 19–21).

While microbial biofilms usually promote survival in diverse environments (7), certain chemicals, including antibiotics, significantly influence biofilm formation and stability (22). Depending on concentration, these chemicals either promote or inhibit biofilm development by interfering with various processes such as initial attachment (23), quorum sensing (24–26), production of extracellular polymeric substance (27), or by directly impacting growth (28, 29).

The establishment of the native gut microbiota in the Western honey bee, *Apis mellifera*, relies on biofilm formation by at least some community members (30). This community is dominated by five core bacterial genera, including the Gram-negatives, *Snodgrassella* and *Gilliamella*, and the Gram-positives, *Lactobacillus* (previously named Firm-5), *Bombilactobacillus* (previously named Firm-4), and *Bifidobacterium* (31). Some additional honey bee-restricted bacteria, including the Gram-negatives *Frischella* and *Bartonella*, are sometimes present (31). *Snodgrassella* forms biofilm directly on the wall of the bee ileum, and mutagenesis analyses showed that genes underlying biofilm formation are required for normal gut colonization (32). Biofilm formation by the bee gut community affects host immune responses and resistance to pathogens (33–35). For example, *Frischella* biofilm in the pylorus region triggers both cellular and humoral immune responses (36). Nonetheless, limited knowledge exists regarding the biofilm-forming potential of other core members, despite some studies showing that all core members can individually colonize the guts of recently emerged bees (35, 37–41).

Among chemicals shown to impair extracellular matrix formation in certain bacterial biofilms is the widely used herbicide glyphosate (42). The main mechanism of action known for glyphosate is the inhibition of the shikimate pathway. This pathway is crucial for the biosynthesis of aromatic metabolites, some of which are known to be important for biofilm formation in certain microbes (43–45). However, whether shikimate pathway inhibition by glyphosate is associated with biofilm disruption is unknown. Glyphosate exposure disrupts the bee gut microbiota and, in particular, reduces abundance of *Snodgrassella* (46–48). Potentially, this impact reflects interference with the ability of *Snodgrassella* to form biofilm within the gut.

In this study, we explore the ability of core honey bee microbiota members to individually colonize the gut and form biofilms in vitro. We also investigate the effects of glyphosate and a glyphosate-based herbicide formulation on the growth and biofilm formation of bee gut symbionts. For comparison, we extend these findings to biofilm-forming bacterial species associated with other animals and plants.

## Results and Discussion

### Core microbiota members independently colonize the bee gut

We first investigated whether specific strains from the bee core microbiota can colonize the gut of recently emerged bees lacking microbiota (Figure 1A). We employed both qPCR and 16S rRNA amplicon sequencing for bees colonized with single strains or a combination of two strains from the same species of core members of the bee gut microbiota. Additionally, we included colonization by the opportunistic bacterium *Serratia marcescens*, by a defined community of native bee gut strains, and by fresh gut homogenates containing the full natural diversity found in bees living in outdoor hives. Based on qPCR data, all individual strains or strain combinations resulted in successful colonization, with high bacterial 16S rRNA transcripts detected in the guts of colonized bees, particularly when compared to control bees that received no treatment (Figure 1B). The average 16S rRNA copy abundance in control bees was about 10^4^, while the abundance in treatments ranged from 10^6^ to 10^11^. By performing 16S rRNA amplicon sequencing on a subset of bees from each cup cage within each group, we determined that most samples were indeed only colonized by the bacterial strains provided in the treatment (Figure 1C). This shows that members of the bee core microbiota can robustly colonize the bee gut in the absence of other community members. Furthermore, defined communities consisting of core member strains as well as gut homogenate treatments allowed typical bee gut colonization by core members, as observed in other studies (49–51). Exceptions included one cup cage from *Snodgrassella*-treated bees, in which considerable *Serratia* reads were detected, suggesting potential cross-contamination, and all *Serratia*-treated bees, in which reads were detected for core and non-core bee gut *Lactobacillus* species and others (Figure 1C).

**Figure 1.**
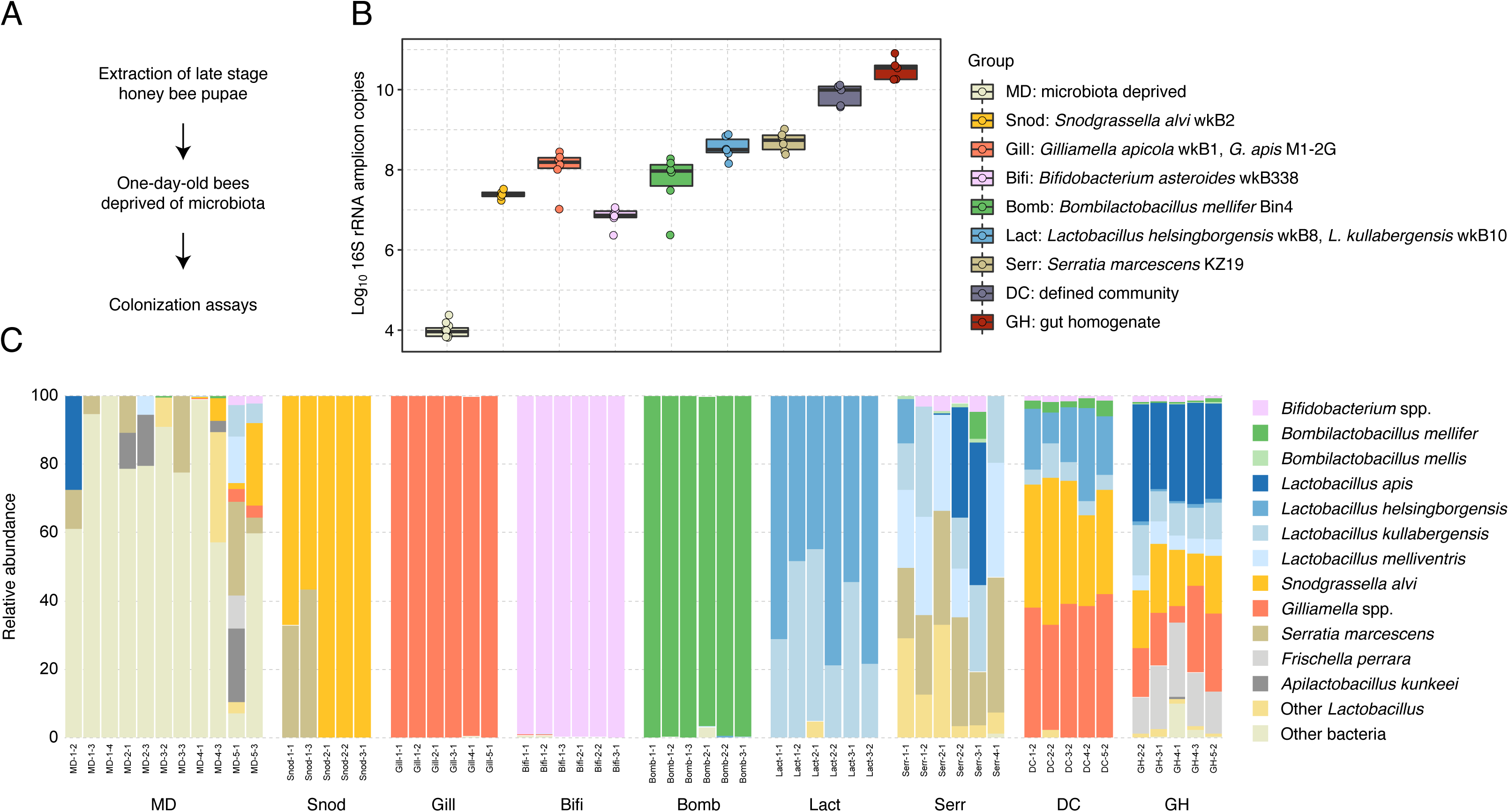
Core members of the honey bee microbiota independently colonize the bee gut. (**A**) Experimental design to obtain microbiota-deprived bees for colonization assays. (**B**) Box plots of absolute abundance of bacteria in the guts of 5-day old honey bees from the colonization assays. (**C**) Stacked column graphs showing the relative abundance of gut bacterial species detected in 5-day old honey bees from the colonization assays. Groups: MD = microbiota-deprived bees; Snod = bees colonized with *Snodgrassella alvi* wkB2; Gill = bees colonized with *Gilliamella apicola* wkB1 and *Gilliamella apis* M1-2G; Bifi = bees colonized with *Bifidobacterium asteroides* wkB338, Bomb = bees colonized with *Bombilactobacillus mellifer* Bin4; Lact = bees colonized with *Lactobacillus helsingborgensis* wkB8 and *Lactobacillus kullabergensis* wkB10; Serr = bees colonized with *Serratia marcescens* KZ19; DC = bees colonized with a defined community of strains wkB2, wkB1, M1-2G, wkB338, Bin4, wkB8 and wkB10; GH = bees colonized with gut homogenates from healthy worker bees from hives.

Several previous studies have colonized newly emerged honey bees, which lack their native microbiota, with single or a set of specific strains of core microbiota members to investigate colonization patterns, their roles in the gut, and host health. Most of these studies confirm colonization status by plating gut homogenates on selective media or by extracting DNA from bee guts and performing quantitative PCR (qPCR) with universal or species-specific 16S rRNA primers (33, 34, 36–40, 50–56). While valid for most purposes, these techniques may not detect potential contamination. 16S rRNA amplicon sequencing enables validation of colonization by each core member and detection of contaminant populations. When reads from other microbes are detected at significant levels, these samples should be either discarded or analyzed cautiously, particularly in studies investigating colonization patterns and impacts of specific microbes on host physiology.

### In vitro biofilm formation is a common feature in bee gut bacteria

One way microbiota members establish themselves in the gut is through biofilm formation or by exploiting the presence of biofilm-producing members for colonization. Specific members of the honey bee gut microbiota, such as *Snodgrassella* (a core member) and *Frischella*, are known for robust biofilm formation in the ileum and pylorus, respectively (32, 57). However, there is limited knowledge about the biofilm-forming potential of other core members, such as *Gilliamella*, *Bifidobacterium*, *Bombilactobacillus* and *Lactobacillus* species. Given that these bacteria can colonize as single strains in the bee gut, we next investigated biofilm-forming capabilities in some strains, as well as in opportunistic bacteria such as *Serratia marcescens*.

Bacterial strains exhibited variable in vitro growth, with *Snodgrassella* and *Gilliamella* strains growing more slowly than *Bifidobacterium*, *Bombilactobacillus*, *Lactobacillus*, and *Serratia* strains (Figure 2A). While all *Snodgrassella* strains formed biofilms in vitro, consistent with previous studies (32), the other members showed variable biofilm formation, with only specific strains displaying biofilm-forming potential (Figure 2B). *Snodgrassella*, *Gilliamella*, *Bifidobacterium*, and *Serratia* strains were cultured in the same media (InsectaGro), but they exhibited no clear correlation between growth rates (measured as OD600) and biofilm formation (measured as OD550). For example, *Snodgrassella* strains had lower growth rates than did *Bifidobacterium* and *Serratia* strains but formed stronger biofilms. *Bombilactobacillus* and *Lactobacillus* required an alternate medium (MRS) for growth. Their higher growth rates may have contributed to robust biofilm formation by specific strains (Figure 2). These findings underscore the potential of the bee core microbiota to form in vitro biofilms, possibly influencing their independent colonization of the bee gut, as seen in vivo.

**Figure 2.**
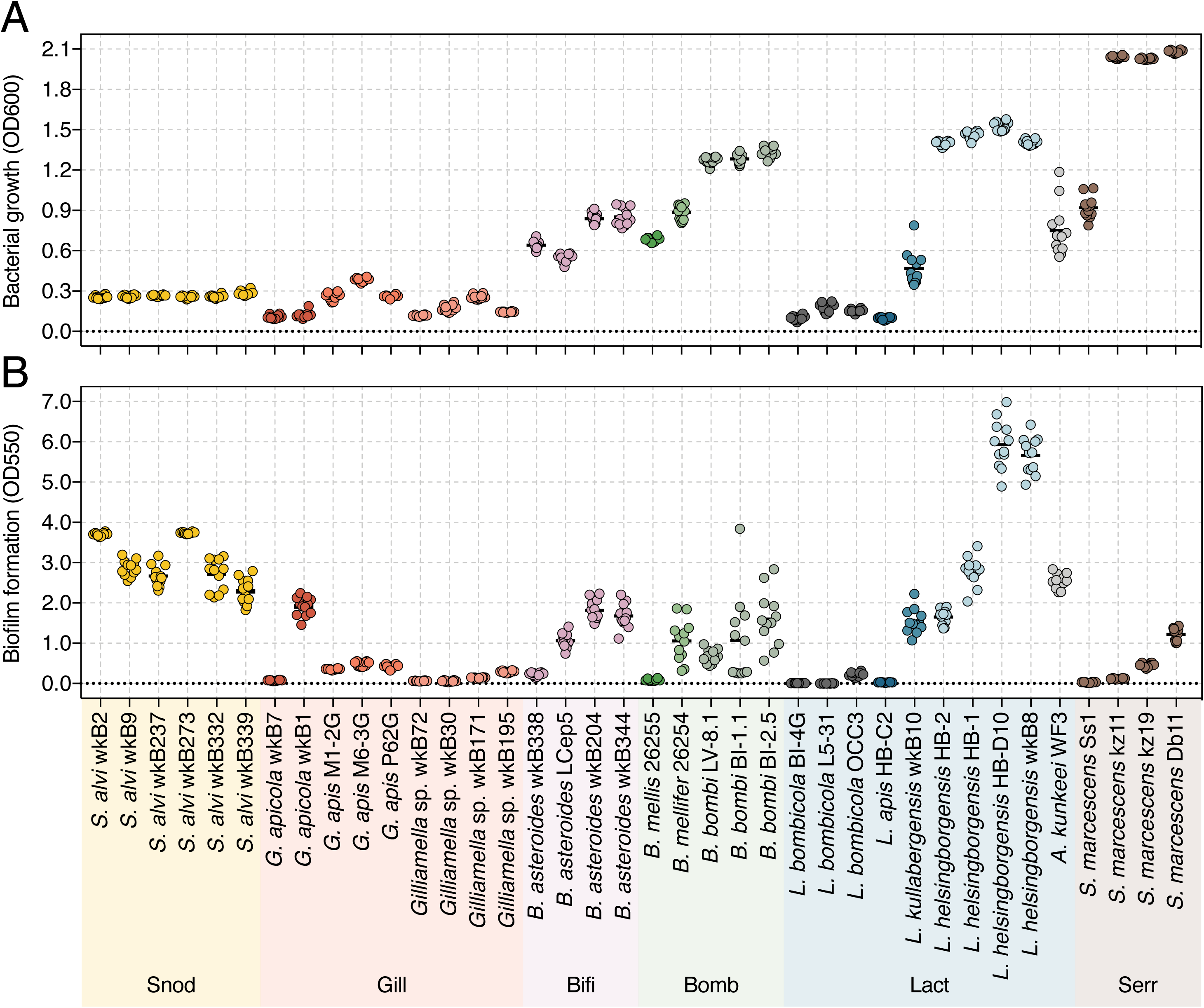
In vitro growth and biofilm formation abilities of honey bee gut symbionts. (**A**) Bacterial growth was measured as optical density at 600 nm after 2 days of incubation at 35 °C and 5% CO_2_. (**B**) Biofilm formation was measured as optical density at 550 nm after washing of planktonic cells, staining attached cells with crystal violet, and adding acetic acid.

Particularly intriguing is the divergent biofilm formation potential of *Serratia marcescens* strains. These strains exhibited markedly greater in vitro growth than bee gut commensals yet showed relatively limited biofilm-forming capabilities (Figure 2). While these in vitro conditions do not mimic the bee gut environment, previous results demonstrate that an intact microbiota can hinder *S. marcescens* proliferation within the bee gut (35). Unlike *S. marcescens* strains KZ11 and KZ19, isolated from bee guts (58), *S. marcescens* strain Ss1, isolated from bee hemolymph (59), exhibited no capacity for in vitro biofilm formation under tested conditions. Additionally, the *S. marcescens* strain Db11, associated with *Drosophila* (60), demonstrated in vitro biofilm formation potential, and is known to be pathogenic to honey bees (58).

### Glyphosate impacts bee gut bacterial growth and biofilm formation

Our next step was to explore how biofilm formation by bee gut bacteria is affected by glyphosate, an herbicide with bacteriostatic properties, and by a glyphosate-based commercial herbicide formulation.

Glyphosate inhibits the shikimate pathway, which is absent from animals but present in plants and several bacteria. This pathway leads to the production of crucial aromatic metabolites, such as aromatic amino acids and certain secondary metabolites (61). Inhibition by glyphosate often impedes growth and leads to death. The target of glyphosate is the enzyme 5-enolpyruvyl-shikimate-3-phosphate synthase (EPSPS). Prior research indicates that bee gut microbiota members differ in whether they encode a Class I susceptible (e.g., *Snodgrassella*, *Gilliamella*, and *Bifidobacterium* strains) or Class II tolerant (e.g., *Bombilactobacillus*) version of EPSPS (48) (Figure 3A and Figure S1). Some members lack EPSPS (e.g., honey bee-restricted *Lactobacillus*) (48) (Figure 3A).

**Figure 3.**
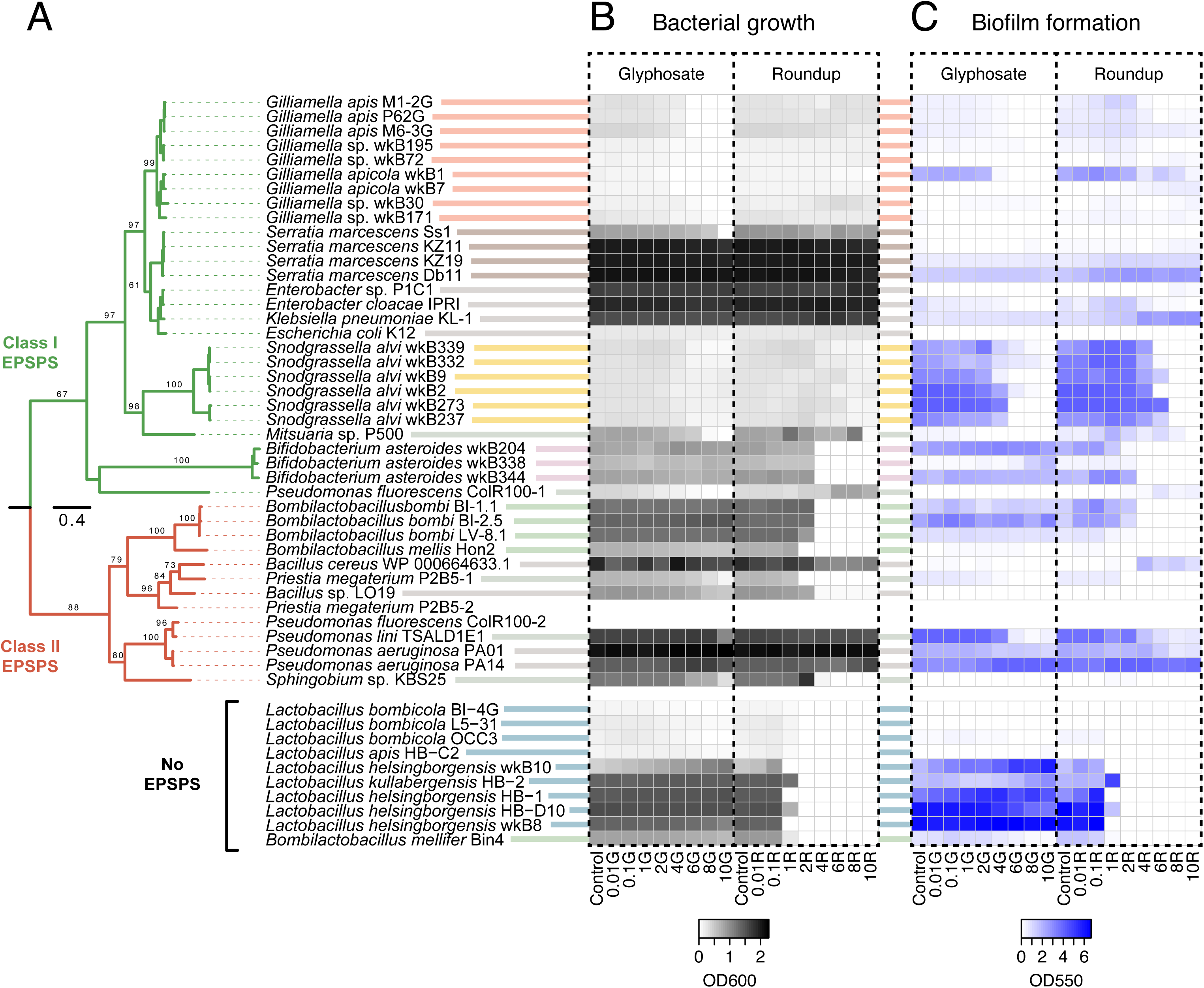
Impacts of glyphosate and a glyphosate-based formulation (Roundup PROMAX®) on growth and biofilm formation of honey bee symbionts and other diverse bacteria. **(A)** The phylogenetic tree was constructed based on the amino acid sequences of 5-enolpyruvylshikimate-3-phosphate synthase (EPSPS) from tested strains with available genomic information (PhyML 3.1, LG model + Gamma4, 100 bootstrap replicates). The heatmap displays the growth (**B**) and biofilm formation (**C**) of bacterial strains in the presence of varying concentrations of glyphosate, either pure or in formulation (ranging from 0.01 to 10 mM). Strain names and sources are listed in Table S7. The genomes of strains P2B5 and ColR100 contain two genes encoding EPSPS, designated as versions “-1” and “-2” in the phylogeny. Since the growth and biofilm data for both versions originate from the same strain, we only included the data once next to version “-1” in the heatmap.

To assess the impact of glyphosate on biofilm, bee gut bacterial strains capable of in vitro biofilm formation were cultured with varying glyphosate concentrations, ranging from 0.01 mM to 10 mM (1.69–1690 ppm). These concentrations simulate different environmental scenarios: minimal exposure, commonly found in water sources (62–65); moderate concentrations detected in nectar (0.02–0.23 mM or 2.78–31.3 ppm) and pollen (0.63–4.57 mM or 87.2–629 ppm) collected from honey bees foraging on plants recently sprayed with glyphosate-based formulations in semi-field experiments (66); and higher, though still below worst-case, concentrations for topical exposure (67, 68). The high concentrations were also used to counteract in vitro culture conditions with abundant nutrients, including some produced by the shikimate pathway, that could mask inhibition by glyphosate.

Glyphosate or a glyphosate-based formulation negatively impacted the growth of Gram-negative core bacteria *Snodgrassella* and *Gilliamella* in a dose-dependent manner, correlating with biofilm inhibition (Figure 3, Figure S2, Figure S3). While some *Snodgrassella* strains, such as wkB2, were less affected by high doses of glyphosate, biofilm formation was nonetheless strongly inhibited (Figure S2). The effects of glyphosate on Gram-positive core bacteria were more variable. While several strains did not experience changes in their growth rates due to glyphosate, others were negatively affected, and some even showed an increase in their growth rates when exposed to glyphosate (Figure 3, Figure S4, Figure S5, Figure S6). For instance, *Lactobacillus helsingborgensis* wkB8 exhibited dose-dependent increases in growth rate and biofilm formation (Figure 3, Figure S6). Among strains that did not exhibit changes in growth rates, glyphosate had diverse effects on biofilm formation. Specific strains displayed dose-dependent biofilm enhancement (e.g., *Bifidobacterium asteroides* wkB204 and wkB338 (Figure 3, Figure S4), as well as *Lactobacillus helsingborgensis* HB-1 (Figure 3, Figure S6)), while others showed dose-dependent biofilm inhibition (e.g., *Lactobacillus helsingborgensis* HB-D10 (Figure S6)). However, most Gram-positive strains were significantly impacted by the herbicide formulation (Figure 3, Figure S4, Figure S5, Figure S6), suggesting potential involvement of formulation components that either enhance glyphosate’s action or exert independent effects on these microbes.

This strain-level variation makes it challenging to predict the impact of glyphosate on bee gut microbiomes, which exhibit strain diversity. This diversity may explain differences in results among published studies (46–48, 67, 69–75). For example, *Bombilactobacillus* and *Lactobacillus* abundances have sometimes decreased or increased in bee guts after glyphosate exposure (48, 70). A similar situation holds for *Gilliamella* (46, 47), while impacts on *Snodgrassella* abundance are consistently negative and are dose-dependent (47).

Glyphosate negatively impacted the growth of *S. marcescens* strains Ss1, KZ11 and KZ19, while the formulation impacted the growth of strains Db11, KZ11 and KZ19 (Figure S7, Supplementary file 1). Although *Serratia* strains encode glyphosate-susceptible EPSPS, glyphosate did not impact biofilm formation, and the glyphosate-based formulation tested even enhanced biofilm formation in a dose-dependent manner for all strains (Figure S7, Supplementary file 2). Therefore, these *S. marcescens* strains may not rely on the shikimate pathway for biofilm formation or may be able to uptake all components required for biofilm formation from the media. Another possibility is that these strains have acquired mutations in EPSPS, conferring resistance to glyphosate. However, no alterations were observed in their conserved amino acid sites to which glyphosate binds (Figure S8) (76, 77). Whether changes in non-conserved sites may contribute to glyphosate resistance deserves further investigation.

### Glyphosate impacts Snodgrassella alvi wkB2 proteome profile

We then decided to investigate why *S. alvi* wkB2, despite possessing a glyphosate-susceptible version of EPSPS (48), can still grow in the presence of high doses of glyphosate but does not form a biofilm. To this end, we cultivated wkB2 in 96-well plates, both in the absence and presence of 10 mM glyphosate (Figure 4A), to investigate growth, biofilm formation, cell viability in the biofilm and supernatant, as well as proteome profiles of whole cultures.

**Figure 4.**
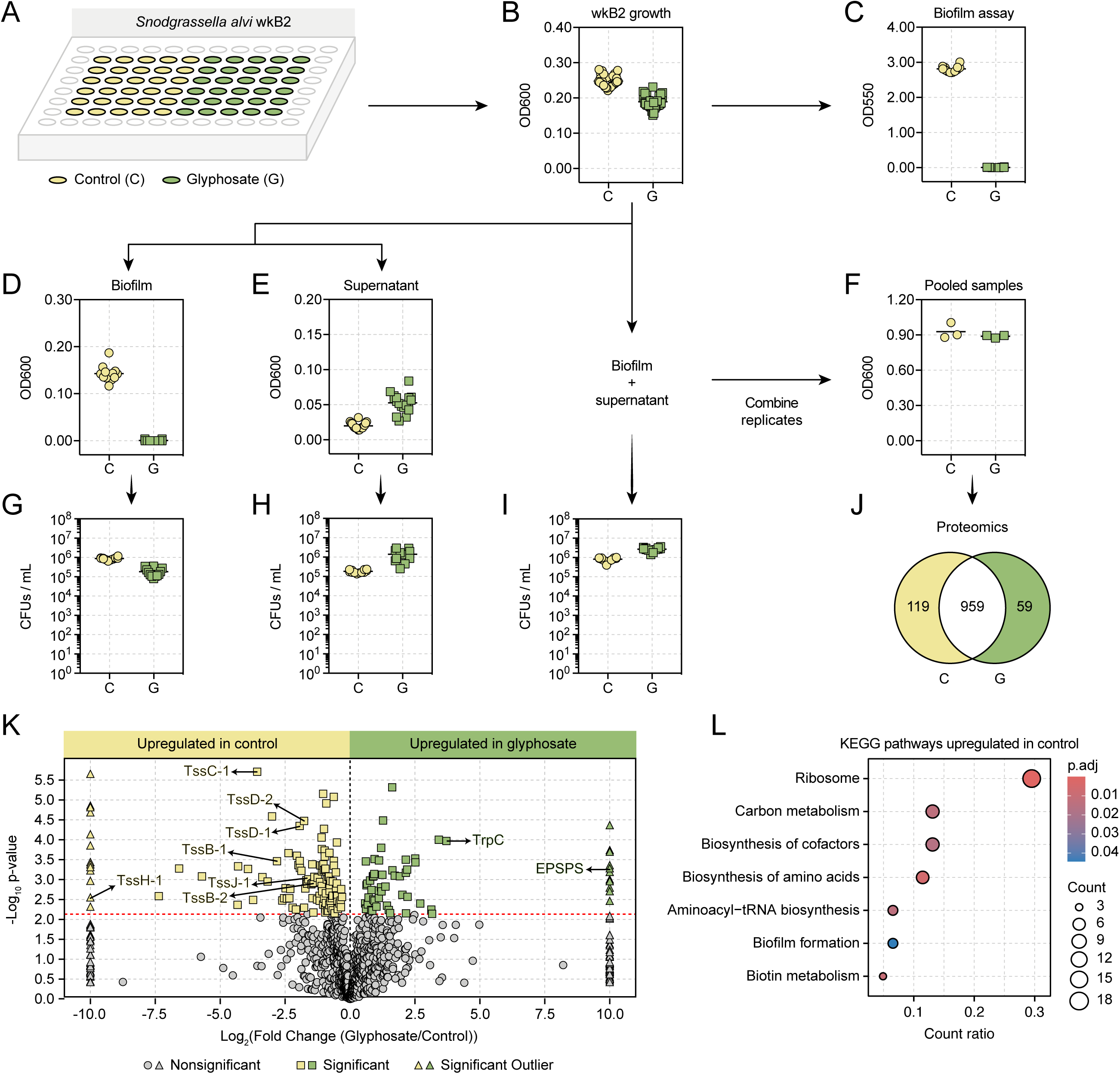
Impact of glyphosate on *Snodgrassella alvi* wkB2 growth, biofilm formation, and proteome. (**A**) Experimental design in in 96-well plates. (**B**) Growth as OD600 measurement in a plate reader after 2 days. (**C**) Biofilm formation as OD550 measurement in a plate reader after performing the crystal violet assay in 2-day old cultures (n = 12 per group). (**D**) OD600 measurements for biofilm samples separated from supernatant and resuspended in PBS solution, and (**E**) OD600 measurements for respective supernatants (n = 18 per group). (**F**) OD600 measurements for biofilms resuspended in their own supernatants and pooled in groups of 6 replicates (n = 3 per group). (**G**) CFU counts in biofilm only and (**H**) supernatant only (n = 12 per group). (**I)** CFUs counts in biofilm resuspended in supernatant samples (n = 12 per group). (**J**) Venn diagram and (**K**) volcano plot showing the number of differentially expressed proteins in control and glyphosate-treated wkB2 cultures (P < 0.05, t-test followed by Benjamini-Hochberg procedure to control for false discovery rate). (**L**) KEGG pathways upregulated in the control group when compared to the glyphosate-treated group based on differentially detected proteins.

As in other trials, wkB2 was able to grow in both conditions, with an average OD600 of 0.25 in control samples and 0.19 in glyphosate-treated samples (Figure 4B), and biofilm formation was only observed in control samples (Figure 4C). Upon separating the supernatant from the biofilm, the OD600 and cell viability were higher in the washed and resuspended biofilms of control samples compared to glyphosate-treated samples (Figure 4D-G). However, an opposite trend was observed for the supernatant, with control samples having lower OD600 and cell viability than glyphosate-treated samples Figure E-H).

Since biofilm is not formed in the presence of glyphosate, we decided to use both biofilm and supernatant for downstream analysis. Therefore, some replicates were homogenized (i.e., biofilm, when present, was resuspended in the supernatant) and pooled in groups of 6 replicates per final replicate, totaling 3 final control samples and 3 final glyphosate-treated samples (Figure 4F). Surprisingly, cell viability was lower in control samples than in glyphosate-treated samples, suggesting that dead cells in the biofilm contribute to the higher OD600 observed for control samples (Figure 4I). OD600 was adjusted to compensate for this, and pooled samples were submitted for proteomics to investigate the mechanism by which wkB2 tolerates glyphosate and the mechanism by which glyphosate potentially impacts biofilm formation.

A total of 1137 proteins were detected in the proteomics analysis, with 59 upregulated proteins in glyphosate-treated samples and 119 upregulated proteins in control samples (Figure 4J and Table S1). In glyphosate-treated samples, KEGG enrichment pathway analysis identified proteins associated with the biosynthesis of amino acids and cofactors, as well as alanine, aspartate and glutamate metabolism (Table S2). The most interesting discovery was the upregulation of the enzymes EPSPS, the known target of glyphosate in the shikimate pathway, and indole-3-glycerolphosphate synthase (TrpC), a branchpoint for the biosynthesis of tryptophan (Figure 4K and Figure S9). These changes in gene expression are likely the main mechanisms employed by wkB2 to maintain activity of the shikimate pathway and aromatic amino acid biosynthesis under glyphosate exposure. Moreover, gene ontology (GO) analysis identified the oxidation-reduction process as the main GO biological process upregulated in glyphosate-treated samples (Figure S10, Table S3), suggesting a mechanism to mitigate the harmful effects of oxidative stress caused by glyphosate and maintain cellular homeostasis.

In control samples, KEGG enrichment pathway analysis found upregulation of proteins associated with translation, biosynthesis of amino acids and cofactors, and biofilm formation (Figure 4L, Table S2). Among the biofilm-upregulated proteins were some associated with Type VI Secretion System 1 (T6SS-1) in *S. alvi*, which exhibits homology to one of the T6SSs in *P. aeruginosa* (78), and includes TssB-1, TssC-1, TssD-1 and TssH-1 (Figure 4K, Figure S11, Table S4). Since the wkB2 genome encodes for two T6SSs (78), we manually inspected and found that some proteins from T6SS-2 were also upregulated (TssB-2 and TssD-2). We also manually inspected our data to investigate the presence of other proteins associated with biofilm formation in wkB2, such as Type IV pilus and some trimeric autotransporter adhesins (*staA* and *staB*) (32, 79, 80). These proteins were indeed detected but showed no significant differences between the control and glyphosate-treated groups (Table S1, Table S5).

T6SSs are complex multiprotein structures used by certain Gram-negative bacteria to deliver toxic effector proteins directly into neighboring cells, facilitating competition or survival in their environment. While T6SSs can confer a competitive advantage to bacteria, they are also a resource-intensive system for bacteria to maintain and operate. These costs may impact bacterial survival under stressed conditions. Potentially, wkB2 can respond to glyphosate-induced stress by reducing production of T6SS-associated proteins. T6SS-associated proteins were the only biofilm-related proteins more abundant in control samples than in glyphosate-treated samples, raising the possibility that T6SS systems may contribute to biofilm formation in *S. alvi* wkB2.

### Glyphosate impacts biofilm formation in other animal- and plant-associated bacteria

To more comprehensively examine the effect of glyphosate on bacterial biofilms, we expanded our in vitro assays to include other animal- and rhizosphere-associated bacteria (Table 2), such as *Pseudomonas* strains, which are known to have biofilm-forming capabilities with host and environmental impacts (81). These strains were cultured under similar conditions as for *Snodgrassella* and *Gilliamella* strains (i.e., InsectaGro) to enable potential comparison (Figure S12).

Our in vitro assays revealed that glyphosate or the tested glyphosate-based formulation displayed a wide range of impacts on the growth of animal-associated strains, ranging from negative to neutral to positive. For example, while glyphosate or the formulation did not impact the growth of *P. aeruginosa* PA01, both negatively impacted biofilm formation (see Figure 3 and Figure S13). On the other hand, both glyphosate and the formulation positively impacted the growth and biofilm formation of *P. aeruginosa* PA14 (Figure 3 and Figure S13). These strains are opportunistic pathogens for humans and are commonly used in the laboratory for studying the pathogenesis of *P. aeruginosa* infections (19). They possess a glyphosate-resistant version of EPSPS (Figure S1), indicating that the observed effects on growth and/or biofilm are linked to other physiological impacts.

Regarding rhizosphere-associated bacteria, two *Pseudomonas* strains (TSALD1E1 and ColR100), *Bacillus* sp. P2B5, *Mitsuaria* sp. P500, and *Streptomyces* sp. P21 were able to form biofilms in vitro (Figure 3, Figure S12, Figure S14). *Pseudomonas lini* TSALD1E1 growth was not impacted by glyphosate concentrations up to 8 mM, but impact on biofilm formation started with lower concentrations of glyphosate, suggesting that the shikimate pathway may play a role in biofilm formation for this strain. Although TSALD1E1 encodes a glyphosate-resistance EPSPS (Figure S1), its growth is still inhibited by 10 mM glyphosate. Conversely, glyphosate negatively impacted *Pseudomonas fluorescens* ColR100 growth at lower doses, yet the formulation had a positive, dose-dependent influence. This strain did not form as robust a biofilm as TSALD1E1, at least under the tested conditions, although high doses of the formulation promoted biofilm formation. Interestingly, ColR100 harbors two versions of EPSPS, glyphosate-susceptible and glyphosate-resistant (Figure S1), yet its growth is still inhibited by lower doses of glyphosate, suggesting a lack of function or expression of the glyphosate-resistant version of EPSPS. Regarding the other strains, glyphosate or the formulation had varying negative impacts on growth and biofilm formation, with stronger effects from the formulation.

## Conclusions

Our results show that glyphosate and a glyphosate-based formulation have variable impacts on growth and biofilm formation depending on the bacterial species and strain. These impacts may be associated with the ability of glyphosate to inhibit EPSPS, but they may reflect other mechanisms that deserve further investigation. In the case of the honey bee symbiont *S. alvi* wkB2, we observed that glyphosate exposure prevents biofilm formation but causes a sharp increase in production of EPSPS. As the wkB2 EPSPS is itself sensitive to glyphosate inhibition (48), this upregulation in the presence of glyphosate is likely mechanism through which this strain achieves tolerance to glyphosate. For several of the other tested strains in which glyphosate inhibited or promoted biofilm formation, the mechanisms may be completely different. The consequences of such impacts will depend on the roles of the exposed microbes in their environment or in a host.

## Material and Methods

### In vivo assays

Pupae of *Apis mellifera* workers in their late developmental stage, characterized by pigmented eyes and limited mobility, were extracted from brood frames within a hive located at the University of Texas at Austin. These pupae were then placed on Kimwipes in sterile plastic containers and kept at 35 °C and around 60% relative humidity to replicate hive conditions. After a few days, approximately 400 newly emerged workers, which naturally lack their native microbiota (82), were randomly divided into nine treatment groups, each with 3 to 5 cup cages, and each cage housing five bees. Group 1 served as the control group and received sterile pollen enriched with a mixture of equal parts sterile 1x PBS and sterile sucrose syrup, maintaining the bees as microbiota deprived. Groups 2 to 7 were exposed to specific bee gut symbionts (Table S6). Bacterial strains were cultured in InsectaGro (for groups 2, 3, 4, and 7) or MRS broth (for groups 5 and 6) at 35°C and 5% CO_2_ overnight. Bacterial cultures were adjusted to an optical density (OD600) of 1, washed with 1x PBS, and diluted in equal parts sterile 1x PBS and sterile sucrose syrup. Each cage in these groups received 200 μL of the respective bacterial suspension mixed with sterile pollen. Group 8 was exposed to a defined community consisting of all bacterial strains provided to groups 2 to 6. This community was created by combining equal portions of each bacterial culture, each adjusted to OD600 of 1. Each cage in this group received 200 μL of this defined community mixed with sterile pollen. Group 9 acquired the native microbiota by being exposed to a fresh bee gut homogenate mixed with sterile pollen. The gut homogenate was prepared by extracting the guts from ten healthy workers from the same hive, mixing them with equal parts 1x PBS and sterile sucrose syrup (5 mL total volume), and transferring 200 μL of this gut homogenate mixed with pollen to each cage. Dead bees were removed daily for five days, and the remaining bees were sampled and stored at -80°C until further analysis.

### RNA extraction and cDNA synthesis

After a 5-day period of exposure, honey bees were sampled from each cup cage, totaling 56 bees (comprising 11 from the MD group, 5 from each of the Snod, DC, and GH groups, and 6 from each of the Gill, Bifi, Bomb, Lact, and Serr groups). RNA extraction was carried out individually for each bee using the Quick-RNA^TM^ Miniprep kit (Zymo Research®). In brief, bee abdomens were crushed in 100 μL of RNA Lysis Buffer, resuspended in 600 μL of the same solution, and transferred to a capped vial along with approximately 0.5 mL of 0.1 mm Zirconia beads (BioSpec Products Inc.). After bead-beating the samples for two rounds of 30 seconds each, they were centrifuged at 14,000 rpm for 30 seconds, and the supernatant was transferred to a new microtube. Subsequent RNA extraction steps followed the protocol provided by Zymo Research®. The resulting RNA samples were eluted in 100 μL of water and stored at -80 °C. RNA quality was assessed through agarose gel electrophoresis and quantified using a NanoDrop^TM^ Lite Spectrophotometer (Thermo Fisher Scientific Inc., USA).

Complementary DNA (cDNA) was synthesized from 1 μg of RNA from each sample using the qScript cDNA Synthesis Kit (QuantaBio, USA) following the manufacturer’s instructions, resulting in a final volume of 20 μL per sample. The cDNA samples were stored at - 20°C.

### qPCR analysis

The cDNA samples (56 in total) were 10-fold diluted for qPCR analysis. Triplicate reactions were set up in 384-well plates on a Thermo Fisher QuantStudio 5 instrument. Each reaction consisted of 5 μL iTaq Universal SYBR Green Supermix (Bio-Rad Inc.), 0.05 μL of 100 μM forward and reverse primers (27F: 5’-agagtttgatcctggctcag-3’ and 355R: 5’- ctgctgcctcccgtaggagt-3’), 3.9 μL H_2_O, and 1.0 μL template cDNA. Cycling conditions included an initial cycle at 50 °C for 2 min and 95 °C for 2 min, followed by 40 cycles at 95 °C for 15 s and 60 °C for 1 min.

Total bacterial 16S rRNA gene transcripts were quantified using standard curves generated by amplifying a conserved 16S rRNA gene region (348 bp) using primers 27F and 355R. The amplicon was ligated into the pGEM-T Easy vector (Promega, USA), which was then transformed into *E. coli* strain DH5-alpha. The vector was purified, digested with ApaI, quantified, and adjusted to 10^10^ copies/μL. Serial dilutions ranging from 10^8^ to 10^2^ copies/μL were prepared as standards, and standard curves were generated. The number of 16S rRNA gene transcripts in each sample was calculated as 10^(Ct-b)/m^, where “b” and “m” are the y-intercept and the slope of the standard curve, respectively. Copy numbers were corrected for the dilution factor.

### 16S rRNA amplicon sequencing and analysis

Diluted cDNA samples (10-fold dilution) served as templates for 16S rRNA library preparation, involving two PCR reactions. PCR 1 amplified the V4 region of the 16S rRNA gene in 25 μL single reactions, including 1 μL of 10 μM forward and reverse primers (Hyb515F: 5’- tcgtcggcagcgtcagatgtgtataagagacaggtgycagcmgccgcggta-3’ and Hyb806R: 5’- gtctcgtgggctcggagatgtgtataagagacagggactachvgggtwtctaat-3’), 12.5 μL of 2× AccuStart^TM^ II PCR SuperMix (Quantabio, USA), and 1 μL of template cDNA. Cycling conditions were 94 °C for 3 min; 30 cycles of 94 °C for 20 s, 50 °C for 15 s, 72 °C for 30 s; followed by 72 °C for 10 min. PCR 2 attached dual indices and Illumina sequencing adapters to PCR 1 products in 25 μL single reactions, including a unique combination of 2 μL of 5 μM index primers (Hyb-F*nn*-i5: 5’- aatgatacggcgaccaccgagatctacacnnnnnntcgtcggcagcgtc-3’, and Hyb-R*nn*-i7: 5’- caagcagaagacggcatacgagatnnnnnngtctcgtgggctcgg-3’), 12.5 μL of 2× AccuStart^TM^ II PCR SuperMix, and 5 μL of PCR 1 product. Cycling conditions were 94 °C for 3 min; 10 cycles of 94 °C for 20 s, 50 °C for 15 s, 72 °C for 60 s; followed by 72 °C for 10 min.

Both PCR product sets were purified with 0.8× HighPrep^TM^ PCR magnetic beads (MagBio, USA) and quantified using a NanoDrop^TM^ Lite Spectrophotometer (Thermo Fisher Scientific Inc., USA). Samples (200 ηg each) were pooled and diluted to a final concentration of 50 ρM. A mixture of 90 μL of the final library and 10 μL of 100 ρM PhiX was loaded onto an Illumina iSeq cartridge, following the manufacturer’s instructions, and subjected to Illumina sequencing on the iSeq platform (2 × 150 sequencing run, instrument model number: FS10000184).

Illumina sequence reads were demultiplexed based on barcode sequences using iSeq software and processed in QIIME 2 version 2019.10 (83). Forward reads were used for downstream analyses due to insufficient overlap and low quality of reverse reads. Primer sequences were removed using the cutadapt plugin (84), reads were truncated to a length of 130 bp, filtered, denoised, and chimeric reads were removed using the DADA2 plugin (85). Taxonomy was assigned to amplicon sequence variants (ASVs) using the SILVA database in the feature-classifier plugin (86). BLASTn searches against the NCBI database were conducted when necessary. Reads with < 1% abundance were removed, along with unassigned, mitochondrial, and chloroplast reads, using the feature-table and taxa filter-table plugins. An Amplified Sequence Variant (ASV) table was generated for investigating microbial composition in colonized bees.

### Preparation of treatments for in vitro assays

Glyphosate standard in its acid form (Caisson Laboratories Inc, lot number: 30322811) was used to prepare stock solutions at 10 mM in Insectagro® DS2 (Corning Inc, lot number: 11622010) or Difco Lactobacilli MRS broth (BD Inc, lot number: 9211338). Working solutions were subsequently prepared with final concentrations of 0.01, 0.1, 1, 2, 4, 6, 8, and 10 mM glyphosate in Insectagro or MRS broth.

Roundup PROMAX® formulation, containing 48.7% (w/v) or 660 g/L of glyphosate potassium salt (equivalent to 540 g/L or 3.2 M of glyphosate acid), was purchased from an agricultural retailer. This formulation was directly diluted in Insectagro or MRS broth, and the experiment’s final concentrations accounted for the initial glyphosate acid concentration in the formulation, referred to here as 0.01, 0.1, 1, 2, 4, 6, 8, and 10 mM Roundup.

### Preparation of bacterial suspensions for in vitro assays

Bacterial strains used in the in vitro assays are detailed in Table S7. All strains were initially cultured in Heart Infusion Agar (Criterion Inc, lot number: 491030) supplemented with 5% Defibrinated Sheep Blood (HemoStat Laboratories Inc, lot number: 663895-2) at 35 °C and 5% CO_2_ for 1 to 3 days. Subsequently, single colonies were transferred to either Insectagro or MRS broth and cultured at 35 °C and 5% CO_2_ overnight. For *Snodgrassella* and *Gilliamella* strains, an additional step was performed to obtain sufficient bacterial mass for subsequent assays. Specifically, 3 mL of each culture was transferred to another tube containing 2 mL of fresh media and cultured again at 35 °C and 5% CO_2_ overnight.

### Bacterial growth assay

The optical density (OD) of each overnight culture was measured at 600 nm using a BioSpectrometer (Eppendorf), and the cells were diluted to an OD600 of 0.5 in Insectagro or MRS broth. For each bacterial culture, 10 μL aliquots of the starter culture were transferred to six biological replicates in a 96-well plate, each containing 190 μL of Insectagro or MRS with varying concentrations of glyphosate (0, 0.01, 0.1, 1, 2, 4, 6, 8, or 10 mM). Similarly, 10 μL aliquots of the starter culture were transferred to six biological replicates in another 96-well plate, each containing 190 μL of Insectagro or MRS with the same range of glyphosate concentrations but in herbicide formulation. Controls consisted of six biological replicates with 200 μL of Insectagro or MRS. The plates were incubated at 35 °C and 5% CO2, and OD600 was measured using a plate reader (Tecan) after 48 hours.

### Biofilm formation assay

Biofilm formation was assessed following a modified version of the protocol described by (87). After a 48-hour incubation period, the liquid in the wells of each plate was removed by inverting the plate and gently shaking it. Subsequently, each plate was immersed in a small container of water to further eliminate unattached cells and residual media components. This rinsing step was repeated twice.

To stain the biofilms, 200 μL of a 0.1% crystal violet solution in water was added to each well of the 96-well plate. The plate was then left to incubate at room temperature for 10-15 minutes. Following incubation, the plate was washed 3-4 times with water using the same immersion technique. Any excess cells and dye were removed by gently shaking the water out of the plate and blotting it on a stack of paper towels. Each plate was inverted and allowed to air dry for 10-15 minutes in a warm room.

For quantitative analysis, 200 μL of a 30% acetic acid solution in water was added to each well of the plate to dissolve the crystal violet. The plate was once again incubated at room temperature for 10-15 minutes. The absorbance was then measured at 550 nm using a plate reader (Tecan), with 30% acetic acid in water serving as the blank.

In cases where the absorbance readings were affected by the strong absorption of crystal violet, 10-fold dilutions of the solutions were prepared to ensure accurate OD550 measurements. This was applied to the following strains: 26254, HB-1, HB-2, HB-D10, wkB8, and wkB10.

### Sample preparation for proteomics analysis

*Snodgrassella alvi* strain wkB2 was cultured in 5 mL of Insectagro at 35°C and 5% CO_2_ overnight. The OD600 was measured and adjusted to 0.5 with Insectagro. Subsequently, 10 μL aliquots were transferred in 96 biological replicates to 96-well plates, with 48 replicates containing 190 μL of InsectaGro and another 48 replicates containing 190 μL of 10 mM glyphosate in InsectaGro. Blanks included 6 replicates of 200 μL of Insectagro and 6 replicates of 200 μL of 10 mM Glyphosate in InsectaGro. The plates were incubated at 35 °C and 5% CO_2_, and after 48 hours, the OD600 was measured using a plate reader (Tecan).

From each group, 12 replicates were reserved for the biofilm formation assay, and 18 replicates were reserved to separate supernatant from biofilm for OD600 measurement and colony forming unit (CFU) counts. The remaining 18 replicates from each group were individually homogenized in the wells by resuspending biofilm (when formed) in the supernatant using wood toothpicks. After homogenization, 10 μL aliquots of some replicates were reserved for CFU counts, and the remaining volume of replicates was pooled into three final biological samples, each consisting of a pool of six replicates, and transferred to 1.5 mL microcentrifuge tubes. The OD600 was measured again, this time using a BioPhotometer (Eppendorf) rather than a plate reader. From each final sample, the OD600 was adjusted to 1 by diluting each sample to a final volume of 0.8 mL.

For cell lysis and protein extraction, 1.2 mL of a bacterial protein extraction reagent (B-PER) solution was added to each replicate – this solution consists of 10 μL of 1 M MgCl_2_, 20 μL of 0.5 M phenylmethylsulfonyl fluoride (in methanol), and 9970 μL B-PER (Thermo Scientific, catalog number: 78248, lot number: LJ148147A). The samples were then incubated for 15 min at room temperature before being frozen until further processing.

Next, samples were centrifuged at 14,000 rpm for 2 min and the supernatant was separated from cell debris. The supernatant was then diluted in 10 mL of exchange buffer (10% glycerol, 1 mM MgCl_2_, 0.1 M NaCl, 1 mM PMSF, and 25 mM Tris pH 8) and concentrated with centrifugal concentrators (10 kDa MWCO, Millipore Sigma-Aldrich, Burlington, MA, USA) to a final volume of 0.1 mL. This process was repeated two more times, after which the final supernatant was transferred to a 1.5 mL vial, and the volume was adjusted to 100 μL with the same exchange buffer.

Subsequently, 30 μL of each concentrated sample was run on a Bolt 4–12% Bis-Tris Plus Gel (Thermo Scientific, catalog number: NW04120BOX, lot number: 21022470). Three final concentrated biological samples from control and glyphosate-treated cultures were then submitted for proteomics analysis at the Biological Mass Spectrometry Facility, University of Texas at Austin. The samples were digested with trypsin, desalted, and run on the Dionex LC and Orbitrap Fusion 2 for LC-MS/MS with a 120 min run time. The data were processed using Proteome Discoverer version 2.5 and Scaffold version 5 (Proteome Software, Inc, Portland, OR, USA). Protein assignment was performed using the NCBI RefSeq amino acid sequence database for wkB2, along with a list of common contaminants, targeting a minimum protein false discovery rate of 0.1%. Upregulated proteins in control and glyphosate-treated samples were used to perform Gene Ontology (GO) analysis using ShinyGo version 0.80 (88). Upregulated proteins were also annotated on the Kyoto Encyclopedia of Genes and Genomes (KEGG) database for pathway enrichment analysis using the R package MicrobiomeProfiler (89).

## Supplementary Figure Legends

**Figure S1.** Maximum-likelihood phylogenetic tree derived from amino acid sequences of 5-enolpyruvylshikimate-3-phosphate synthase (EPSPS). Sequence alignment was performed using Muscle and trimmed to 671 sites, corresponding to amino acids 18 to 423 of the *V. cholerae* sequence. PhyML 3.1 was utilized with the LG model and Gamma4, employing 100 bootstrap replicates. Glyphosate-susceptible Class Iα and Class Iβ, and glyphosate-resistant Class II EPSPS are highlighted in green, gray, and red, respectively.

**Figure S2.** Boxplots show the in vitro growth and biofilm formation abilities of *Snodgrassella alvi* strains wkB2 (A-B), wkB9 (C-D), wkB237 (E-F), wkB273 (G-H), wkB332 (I-J), and wkB339 (K-L) in the presence of variable concentrations of glyphosate, either pure (G) or in herbicide formulation (R), ranging from 0.01 mM to 10 mM. The control consisted of growth in InsectaGro media only (n = 6 per group). Statistical analyses are found in Supplementary Files 1 and 2.

**Figure S3.** Boxplots show the in vitro growth and biofilm formation abilities of *Gilliamella* spp. strains wkB7 (A-B), wkB1 (C-D), M1-2G (E-F), M6-3G (G-H), P62G (I-J), wkB72 (K-L), wkB30 (M-N), wkB171 (O-P), and wkB195 (Q-R) in the presence of variable concentrations of glyphosate, either pure (G) or in herbicide formulation (R), ranging from 0.01 mM to 10 mM. The control consisted of growth in InsectaGro media only (n = 6 per group). Statistical analyses are found in Supplementary Files 1 and 2.

**Figure S4.** Boxplots show the in vitro growth and biofilm formation abilities of *Bifidobacterium* spp. strains wkB338 (A-B), LCep5 (C-D), wkB204 (E-F), and wkB344(G-H) in the presence of variable concentrations of glyphosate, either pure (G) or in herbicide formulation (R), ranging from 0.01 mM to 10 mM. The control consisted of growth in InsectaGro media only (n = 6 per group). Statistical analyses are found in Supplementary Files 1 and 2.

**Figure S5.** Boxplots show the in vitro growth and biofilm formation abilities of *Bombilactobacillus* spp. strains 26255 (A-B), 26254 (C-D), LV-8.1 (E-F), BI-1.1 (G-H), and BI-2.5 in the presence of variable concentrations of glyphosate, either pure (G) or in herbicide formulation (R), ranging from 0.01 mM to 10 mM. The control consisted of growth in InsectaGro media only (n = 6 per group). Statistical analyses are found in Supplementary Files 1 and 2.

**Figure S6.** Boxplots show the in vitro growth and biofilm formation abilities of *Lactobacillus* spp. strains BI-4G (A-B), L5-31 (C-D), OCC3 (E-F), HB-C2 (G-H), wkB10 (I-J), HB-2 (K-L), HB-1 (M-N), HB-D10 (O-P), wkB8 (Q-R), and *Apilactobacillus* sp. WF3 (S-T) in the presence of variable concentrations of glyphosate, either pure (G) or in herbicide formulation (R), ranging from 0.01 mM to 10 mM. The control consisted of growth in InsectaGro media only (n = 6 per group). Statistical analyses are found in Supplementary Files 1 and 2.

**Figure S7.** Boxplots show the in vitro growth and biofilm formation abilities of *Serratia marcescens* strains Ss1 (A-B), KZ11 (C-D), KZ19 (E-F), and Db11 (G-H) in the presence of variable concentrations of glyphosate, either pure (G) or in herbicide formulation (R), ranging from 0.01 mM to 10 mM. The control consisted of growth in InsectaGro media only (n = 6 per group). Statistical analyses are found in Supplementary Files 1 and 2.

**Figure S8**. Detection of conserved amino acid residues characteristic of Class I EPSPS in *Serratia marcescens* strains using the amino acid sequence of *Vibrio cholerae* as a reference. Red boxes highlight conserved amino acid residues.

**Figure S9.** Shikimate pathway and aromatic amino acid biosynthetic enzymes detected in the proteomics analysis comparing control and glyphosate-treated wkB2 cultures. Bar plots show normalized total precursor intensity counts for detected proteins in each group. P < 0.05, t-test followed by Benjamini-Hochberg procedure.

**Figure S10.** Gene Ontology enrichment analysis for control and (A) glyphosate-treated samples (B) using the differentially expressed proteins in each condition.

**Figure S11.** Schematic figures illustrate the proteins that compose Type VI Secretion Systems 1 and 2 in *Snodgrassella alvi* wkB2. Barplots show normalized total precursor intensity counts for detected proteins in each group. P < 0.05, t-test followed by Benjamini-Hochberg procedure.

**Figure S12.** In vitro growth and biofilm formation abilities of diverse animal-associated and rhizosphere-associated bacteria. (A) Bacterial growth was measured as optical density at 600 nm after 2 days of incubation at 35 °C and 5% CO_2_. (B) Biofilm formation was measured as optical density at 550 nm after washing of planktonic cells, staining attached cells with crystal violet, and adding acetic acid.

**Figure S13.** Boxplots show the in vitro growth and biofilm formation abilities of animal-associated bacterial strains in the presence of variable concentrations of glyphosate, either pure (G) or in herbicide formulation (R), ranging from 0.01 mM to 10 mM. The control consisted of growth in InsectaGro media only (n = 6 per group). Statistical analyses are found in Supplementary Files 1 and 2.

**Figure S14.** Boxplots show the in vitro growth and biofilm formation abilities of rhizosphere-associated bacterial strains in the presence of variable concentrations of glyphosate, either pure (G) or in herbicide formulation (R), ranging from 0.01 mM to 10 mM. The control consisted of growth in InsectaGro media only (n = 6 per group). Statistical analyses are found in Supplementary Files 1 and 2.

**Supplementary file 1.** Statistical reports for growth curves assays.

**Supplementary file 2.** Statistical reports for biofilm formation assays.

## Data availability

16S rRNA amplicon sequencing data are available on NCBI BioProject PRJNA1088537. Other data are included in this article and its supplementary information files.

## Acknowledgements

Thanks to the members of the Nancy Moran and Howard Ochman labs, especially Kim Hammond for maintaining hives, Eli Powell for technical assistance, and Paul Kirchberger for providing human or lab-related bacterial strains and for discussion of this project. Thanks to Thomas E. Juenger from the Department of Integrative Biology at the University of Texas at Austin for providing rhizosphere-associated bacterial strains. Thanks to Maria Person, and Michelle Gadush at the UT Austin Center for Biomedical Research Support Biological Mass Spectrometry Facility (RRID:SCR_021728) for processing protein samples.

